# Empirical evaluation of fused EEG-MEG source reconstruction. Application to auditory mismatch generators

**DOI:** 10.1101/765966

**Authors:** Françoise Lecaignard, Olivier Bertrand, Anne Caclin, Jérémie Mattout

**Affiliations:** Lyon Neuroscience Research Center, CRNL; INSERM, U1028; CNRS, UMR5292; Brain Dynamics and Cognition Team, Lyon, F-69000, France; University Lyon 1, Lyon, F-69000, France; CERMEP Imaging Center, Lyon, F-69500, France

**Keywords:** Distributed sources, Multimodal integration, Bayesian model comparison, Group-level inference, Mismatch Negativity (MMN), Early Deviance Response

## Abstract

Since their introduction in the late eighties, Bayesian approaches for neuroimaging have opened the way to new powerful and quantitative analysis of brain data. Here, we apply this statistical framework to evaluate empirically the gain of fused EEG-MEG source reconstruction, compared to unimodal (EEG or MEG) one. Combining EEG and MEG information for source reconstruction has been consistently evidenced to enhance localization performances using simulated data. However, given considerable efforts to conduct simultaneous recordings, empirical evaluation becomes necessary to quantify the real information gain. And this is obviously not straightforward due to the ill-posedness of the inverse problem. Here, we consider Bayesian model comparison to quantify the ability of EEG, MEG and fused (EEG/MEG) inversions of *individual* data to resolve spatial source models. These models consisted in *group-level* cortical distributions inferred from real EEG, MEG and EEG/MEG brain responses. We applied this comparative evaluation to the timely issue of the generators of auditory mismatch responses evoked by unexpected sounds. These included the well-known *Mismatch Negativity* (MMN) but also earlier deviance responses. As expected, fused localization was evidenced to outperform unimodal inversions with larger model separability. The present methodology confirms with real data the theoretical interest of simultaneous EEG/MEG recordings and fused inversion to highly inform (spatially and temporally) source modeling. Precisely, a bilateral fronto-temporal network could be identified for both the MMN and early deviance response. Interestingly, multimodal inversions succeeded in revealing spatio-temporal details of the functional organization within the supratemporal plane that have not been reported so far, nor were visible here with unimodal inversions. The present refined auditory network could serve as priors for auditory modeling studies.

## 1. Introduction

Source reconstruction of electrophysiological responses have become a standard analysis in neuroimaging, as suggested by the increasing number of papers, as well as the numerous methodologies afforded by electrophysiological analysis software. Whatever the methodology (Lecaignard and Mattout, 2015), the ill-posed nature of the underlying inverse problem remains (from a mathematical point of view, recognition of true generators is impossible). This issue calls for highly informed data to be confronted to models, as can be achieved with the integration of EEG and MEG signals proposed more than 30 years ago (Puce and Hamalainen, 2017). This paper addresses the added value of combining EEG and MEG data for distributed source localization, which we evaluated here *empirically* with auditory mismatch responses.

Merging EEG and MEG aims at accounting for information missed by one modality and captured by the other one (Dale and Sereno, 1993; Fuchs et al., 1998), and crucially, at reducing the under-determined nature of the ill-posed inverse problem thanks to complementary information gathered by these two modalities (Plonsey and Heppner, 1967). Fused reconstruction therefore appears promising to reach high temporal and spatial resolutions in brain function imaging. Greater performances for fusion than separate EEG or MEG source reconstructions were indeed consistently reported in simulation-based studies. Quantitative evaluations rested on various metrics obtained from the comparison of the true distribution (that has generated the synthetic data) and reconstructed ones. In short, reduced localization errors could be reported for both superficial and deep sources (Fuchs et al., 1998), as well as for different signal-to-noise ratio (SNR) and sensor montages (Babiloni et al., 2004). Decrease of the undesirable sensitivity of inversion methods to source orientation (Baillet et al., 1999) and enhanced precision of source estimates (Henson et al., 2009b) were also reported. Further evaluation with empirical data is a necessary step, but in this case the ill-posedness of the inverse problem obviously prevents from using simulation-based metrics. To date, only few studies attempted to circumvent this issue. They considered specific cases for which fMRI results (Sharon et al., 2007), widely described median nerve stimulation (Molins et al., 2008) or intracranial recordings with epileptic patients (Chowdhury et al., 2015) were assumed to provide the true solution to be compared with. Noticeably, all these studies were in favor of reduced mislocalizations with fused inversion. In the current study, we propose a general approach for the quantitative evaluation of fusion that applies to any empirical data. We used advanced statistical methods for source reconstruction that formalize model inversion as Bayesian inference (Friston et al., 2006; Mattout et al., 2006). This framework enabled us to exploit Bayesian model comparison (Penny et al., 2004) to investigate the ability of each modality (EEG, MEG and fusion) to separate different source distributions (being spatial models). Our approach thus quantifies the spatial (model) resolution of each modality.

We applied this evaluation to auditory mismatch (or deviance) responses elicited by a change (or deviant) in a regular acoustic environment, including the well-known Mismatch Negativity (MMN) (Näätänen et al., 2007). This choice was motivated by the outstanding place the MMN has occupied in cognitive and clinical neuroscience (Auksztulewicz and Friston, 2016; Morlet and Fischer, 2014; Sussman and Shafer, 2014), contrasting with the arguably poor consistency of findings in the MMN source research (Fulham et al., 2014; Schönwiesner et al., 2007). Beside, recent findings of earlier mismatch responses than the MMN (Escera et al., 2014; Lecaignard et al., 2015) encourage to develop a comprehensive analysis of auditory responses to improve our understanding of auditory (deviance) processing. To date, only a few MEG studies addressed the localization of early deviance components (Recasens et al., 2014a; 2014b; Ruhnau et al., 2013), with activity circumscribed in the primary auditory cortex. Taken together, these recent findings indicate that it is time to combine high temporal and spatial information for an in-depth characterization of auditory processing.

Strong efforts using different neuroimaging techniques have been made to identify the cortical generators of the MMN for about three decades. Functional Magnetic Resonance Imaging (fMRI) and electrophysiological techniques (EEG, MEG) were mostly employed, that favored spatial or temporal precision respectively. To our knowledge no study has been conducted using fused inversion (simultaneous recordings but separate source modeling were conducted in Huotilainen et al., 1998; Kuuluvainen et al., 2014; Rinne et al., 2000). Taken together, fMRI (see for review Deouell, 2007) and electrophysiological studies (Fulham et al., 2014; Giard et al., 1995; Lappe et al., 2013a; Marco-Pallarés et al., 2005; Recasens et al., 2014b; Ruhnau et al., 2013; Waberski et al., 2001) suggested that the most prominent sources are located in temporal and frontal areas. However, there is a large and acknowledged variability across findings that prevents from a reliable and detailed description of the MMN network. It is possible that none of these modalities may be sufficiently informed spatially and temporally when employed alone, which pleads for advanced methods such as fused reconstruction.

In this context, the aim of the current study was twofold: first, to propose a general method to evaluate quantitatively the performance of separate and fused source reconstruction with empirical data. The second aim was to provide a detailed description of early and late auditory mismatch generators using advanced statistical methods including fused inversion (Henson et al., 2009b). We considered data originating from a previous passive auditory oddball study (Lecaignard et al., 2015) with two deviance features (frequency and intensity, separately manipulated) and conducted with simultaneous EEG and MEG recordings. Our results demonstrate the larger spatial model resolution of fused inversion and the usefulness of such information integration that here produced a fine-grained description of a fronto-temporal network underlying auditory processing.

## 2. Material and Methods

This section is divided into three parts: first, we briefly describe the methodologies for source localization employed in the present study including model inversion with group-level inference (Litvak and Friston, 2008) and MEG-EEG fusion (Henson et al., 2009b). Second, we describe our approach for the quantitative evaluation of EEG, MEG and fused MEG-EEG inversions. Finally, the third section presents the multimodal dataset used to validate our approach, resting on simultaneous EEG-MEG recordings of auditory frequency (FRQ) and intensity (INT) deviance responses.

### 2.1. Methods for source reconstruction

#### Forward model computation

For both MEG and EEG modalities, a three-layer realistic Boundary Element Model (BEM) (Hämäläinen and Sarvas, 1989) was employed, with homogenous and isotropic conductivities within each layer set to 0.33, 0.0041 and 0.33 S/m for the scalp, skull and brain, respectively (Rush and Driscoll, 1968). The source domain included *N*_*s*_=20484 sources (mean average distance = 3.4 mm) distributed on the cortical mesh (grey-white matter interface) and we used surface normal constraints for dipole orientation. All meshes derived from canonical uniformly tessellated templates (provided with SPM8) that had been warped from individual MRI to account for subject-specific anatomy (Mattout et al., 2007). Coregistration of the resulting head model and functional data (EEG, MEG) was achieved for both modalities separately using a rigid spatial transformation based on three anatomical landmarks (or fiducials), positioned at nasion, left and right pre-auricular points. For MEG data, head position was averaged across experimental sessions to allow for a common forward model between conditions. For each participant and each modality, computation of accurate BEM was performed with the software Openmeeg (http://openmeeg.github.io) (Gramfort et al., 2010). Re-referencing to the average mastoids was applied to EEG BEM. The resulting lead-field operator or gain-matrix 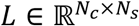 (with *N*_*c*_ sensors and *N*_*s*_ sources) embodying the pre-cited anatomical and biophysical assumptions, enters the following linear generative model *M* of data 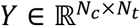 (with *N*_*t*_ time samples):

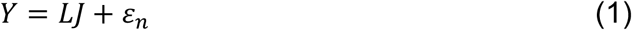

where *J* represents the source distribution, i.e. the magnitude of dipole at each node of the cortical mesh, and *ε*_*n*_ represents the residual error term.

#### Model inversion using Multiple Sparse Priors (MSP)

Within a hierarchical Bayesian framework, we defined *J* as a multivariate Gaussian distribution of the form 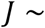 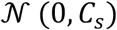 with 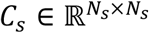 the (unknown) spatial source covariance. We assumed a multivariate Gaussian error term 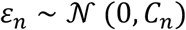 with 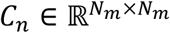 the (unknown) spatial noise covariance (relatively to a normalized spatial space composed of *N*_*m*_ modes that will be defined in the following section). We used Multiple Sparse Priors (Friston et al., 2008b) to estimate both the distribution *J* that satisfies the general equation of linear model with Gaussian errors:

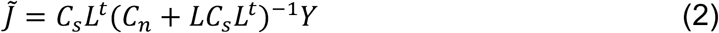

and the posterior distribution of *C*_*s*_ and *C*_*n*_. Precisely, *C*_*s*_ is defined as a linear combination of *N*_*p*_ variance components 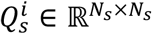 (the sparse priors), weighted by hyperparameters 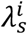:

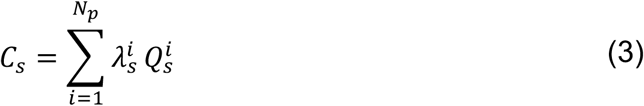

For the initial condition, we used SPM8 default sparse priors including 256 components in each hemisphere, and applied a bilaterality constraint, leading to a total of *N*_*p*_ = 712 variance components. Estimation of the associated hyperparameters 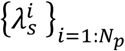 was driven by the principle of source sparsity implemented in the Greedy-Search (GS) algorithm (Friston et al., 2008a). At the sensor level, we assumed a single variance component equal to the identity matrix per modality, with hyperparameter weighting as follows:

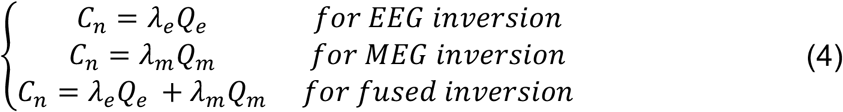

MSP rests upon expectation maximization (EM) and provides Restricted Maximum Likelihood (ReML) estimates of hyperparameters 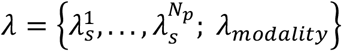 and Maximum A Priori (MAP) estimate of *J* using Eq.(2) (Friston et al., 2007). EM is an iterative process guided by the maximization of the free energy 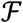, an approximation of the log-evidence of the model (the log-value of *p*(*Y*|*M*), the probability of observing the data *Y* given the generative model *M* defined in Eq.(1)).

#### Group-level inference

Group-level inference (Litvak and Friston, 2008) aims at specifying the prior distribution on the source covariance *C*_*s*_ by accounting for the assumption that distribution *J* should be common to all participants. This is a two-step procedure (*Figure 1*) that we used in the present reconstruction study (using SPM8) and that has also inspired our quantitative evaluation of fused inversion (see below):

- First step performs a single *group-level* inversion using default sparse priors. Resulting posterior hyperparameters are thus informed by the group-level variance of the data; they provide a posterior on *C*_*s*_ (Eq.(3)).
- Second step proceeds to *individual-level* inversions, starting with the group-informed posterior on *C*_*s*_ as prior, here referred to as *group priors*.

**Figure 1.**
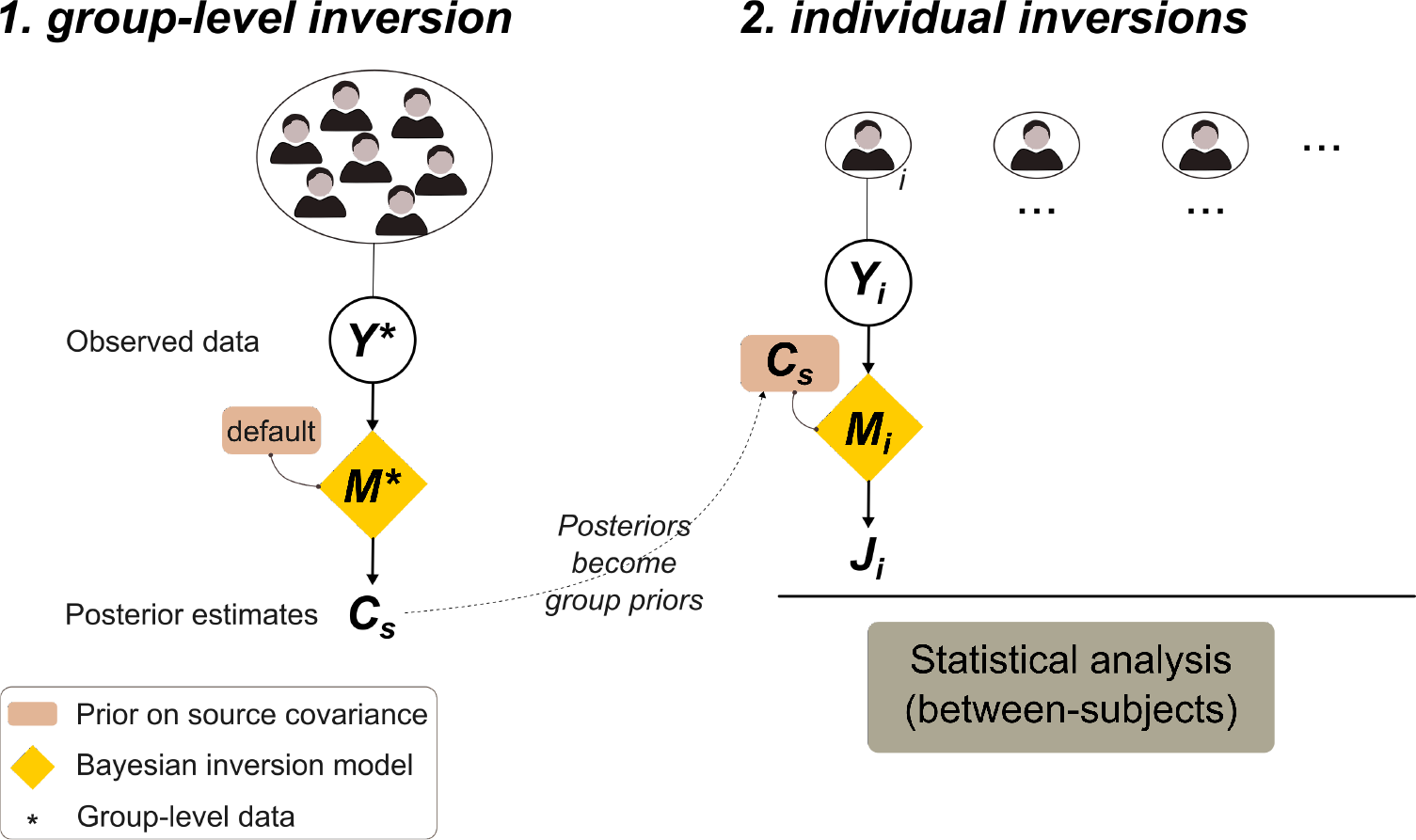
Schematic view of group level inference (Litvak et al. 2008). The two-stage procedure aims at informing subject-specific inversion with source priors deriving from the source distribution common to the group. Notations Y, M, C and J refer to sensor data, inversion model, source covariance and source distribution respectively, as specified in the main text.

In practice, as detailed in Litvak et al., (2008), the second step is left with two/three hyperparameters to estimate: {λ_*s*_; λ_*e*_}, {λ_*s*_; λ_*m*_}, {λ_*s*_; λ_*e*_, λ_*m*_} for EEG, MEG and fused inversions, respectively. Prior to data inversion, group-level inference involves the normalization of the individual sensor-level data in a common spatial-mode space (Friston et al., 2008b). In short, this space is composed of *N*_*m*_ orthogonal virtual sensors (referred to as spatial modes) resulting from the singular value decomposition (SVD) of a group-informed gain matrix. Data reduction is also achieved using a subsequent projection of the data on temporal modes (Friston et al., 2006). For each subject, the spatially and temporally projected data 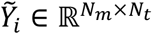 is rescaled (using the trace of 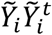) to accommodate signal amplitude differences over spatial modes. After model inversion, the reconstructed source activity *J* is projected on spatial modes and *R*, the percentage of data explained by *J* is computed to quantify the variance explained by *J* relative to the residual variance.

#### Fused MEG-EEG inversion

The fused inversion approach proposed in Henson et al. (2009b) was employed in the current study. This method entails the necessary rescaling of data and gain matrix over modalities to accommodate the different physical nature of signals. This rescaling leads to two crucial aspects: *(1)* projected data on MEG and EEG spatial modes become homogeneous and *(2)* sensor-level hyperparameters λ_*e*_ and λ_*m*_ can be quantitatively compared to assess the relative contribution of each modality to account for the variance of the observed data. Such comparison was conducted using paired Student’s t-tests in the case of the MMN inversion ([150,200] ms) in condition FRQ and INT (see below).

### 2.2. Quantitative evaluation of separate and fused inversions

Bayesian Model Comparison (BMC) is a formal way to quantitatively compare models (*M*_1,_*M*_2_, …), based on their inferred model evidences (*p*(*Y*|*M*_1_), *p*(*Y*|*M*_2_), …) that each quantifies how likely model *M*_*i*_ is to have generated data *Y* (Penny et al., 2010). In the present case, as illustrated in ***Figure 2***, for each modality: EEG (*e*), MEG (*m*), and Fusion (*f*), we conducted a BMC that involved three models differing only on the group priors entering individual inversions. The three variants of group priors were inferred by the group-level inversion of EEG data (*Y*_*e*_), MEG data (*Y*_*m*_), and fused inversion of EEG and MEG data 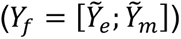. These specific models entail the spatial information that could be captured by each modality over the group of subjects. Our aim was to evaluate the ability of each modality to resolve the resulting source distributions (reconstructed at the group level). Such model separability can be interpreted as a measure of spatial resolution. In the following, for each modality *mod* ∈ {*e*, *m*, *f*}, the three group prior models will be denoted *M*_*mod,e*_, *M*_*mod,m*_ and *M*_*mod,f*_. To run this evaluation, a total of 9 inversions were thus computed for each subject: three modalities for data (*mod*_*d*_, *with d* ∈ {*e*, *m*, *f*}) combined with three modalities for group priors (*mod*_*p*_, *with p* ∈ {*e*, *m*, *f*}). Thereafter, for each modality *mod*_*d*_, the free energy approximating model evidences 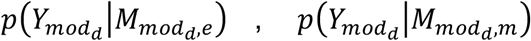 and 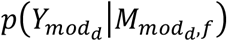 were compared across subjects with BMC using a random effect (RFX) model. To account for inter-individual variability, we also computed the following free energy differences for each subject and for each modality *mod*_*d*_, approximating the log-Bayes Factor:

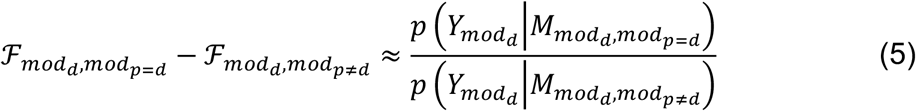

**Figure 2.**
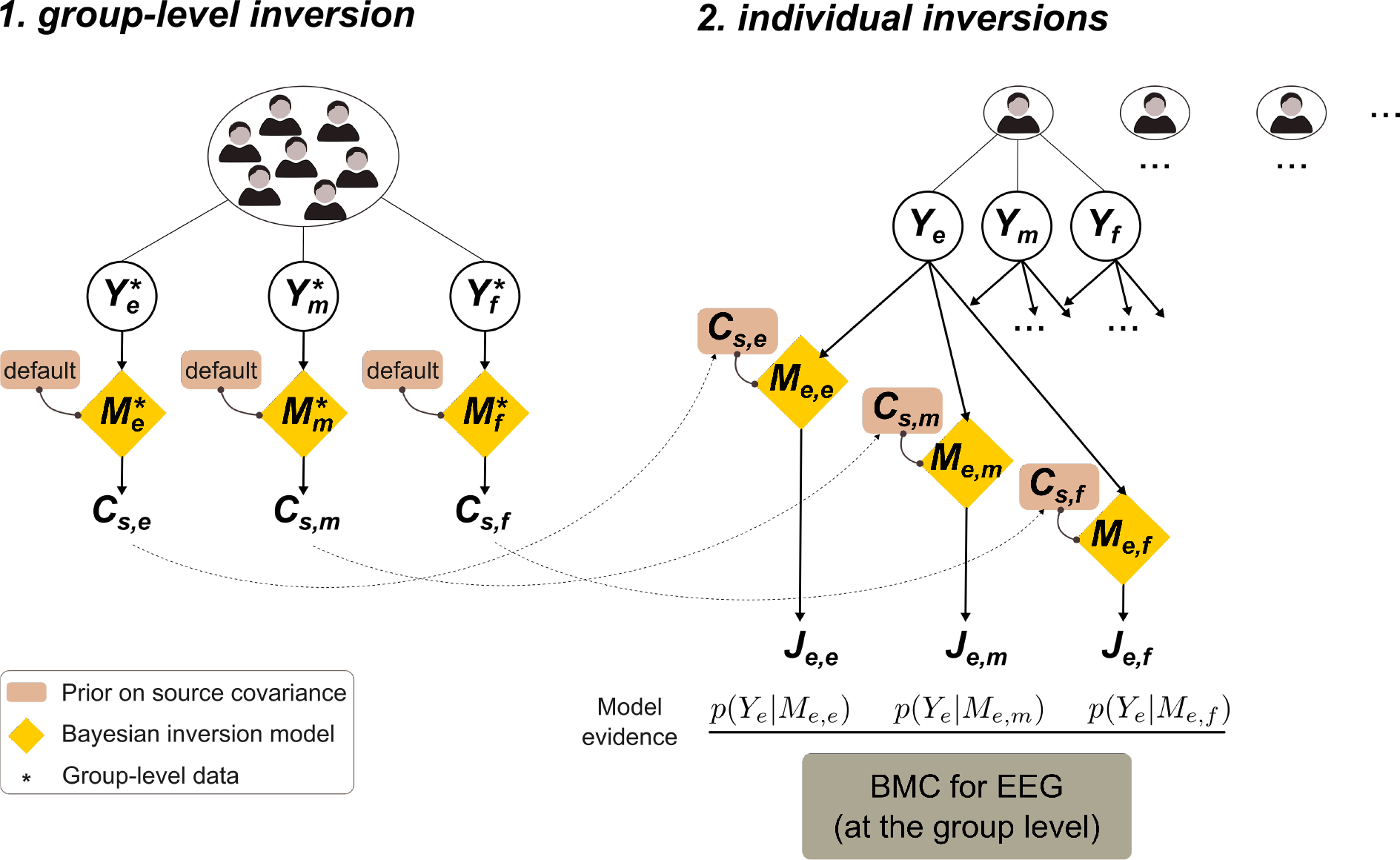
Evaluation scheme for multimodal evaluation. The three separate group-level inversions performed for each modality provides the source priors for subsequent subject-specific inversions (nine per subject). Within each modality (*Y*_*i*_), Bayesian model comparison (BMC) proceeds at the group level using approximated model evidence to select which source prior models (*M*_*i,e*_, *M*_*i,m*_ or *M*_*i,f*_) performs best (example is given in the EEG case in the figure).

Following the usual principles of Kass and Raftery (1995), a free energy difference (in absolute terms) lower than or equal to 3 indicates that models have comparable evidence: related group priors are of equal plausibility. Under the assumption of non-identical group priors across modalities (EEG, MEG and fusion do not capture the same information), we would thus conclude that modality *mod*_*d*_ is not informed enough to disentangle these different models. On the contrary, an absolute difference greater than 3 would support a large spatial resolution of *mod*_*d*_ over model space. We expected *i)* EEG to have a poor capacity to separate group prior models, due to volume conduction which is acknowledged to degrade the spatial resolution of EEG (Vallaghé and Clerc, 2009) and *ii)* Fusion to have the largest resolution, being informed by the complementary EEG and MEG (Lopes da Silva, 2013). An original aspect of the proposed method pertains to the fact that it allows comparing quantitatively EEG, MEG and fused source reconstructions applied to real (not simulated) data. We carried out this empirical evaluation for the frequency and intensity MMN and early deviance response as described below.

### 2.3. Empirical data for source reconstruction and multimodal evaluation

Data originate from a passive auditory oddball study with simultaneous MEG-EEG recordings where the EEG analysis revealed two deviance responses: an early effect occurring within 70 ms after stimulus onset and a late effect (MMN) peaking at 170 ms post-stimulus (Lecaignard et al., 2015). We refer the reader to this study for a more detailed description of material and methods.

#### Participants

27 adults (14 female, mean age 25±4 years, ranging from 18 to 35) participated in this experiment. All participants were free from neurological or psychiatric disorder, and reported normal hearing. All participants gave written informed consent and were paid for their participation. Ethical approval was obtained from the appropriate regional ethics committee on Human Research (CPP Sud-Est IV - 2010-A00301-38). Seven participants were excluded because they paid attention to sounds or their data was of low quality, leading the current analysis based on a total of 20 participants.

#### Experimental design

Oddball sequences embedding frequency and intensity deviants (conditions UF and UI in Lecaignard et al., 2015) were considered in the present analysis, that we rename here as FRQ and INT, respectively. Both sequence types had the same deviant probability (*p* = 0.17). Two different frequencies (*f*_1_=500 Hz and *i*_2_=550 Hz) and two different intensities (*i*_1_=50 dB SL (sensation level) and *i*_2_=60 dB SL) were combined to define the four different stimuli that were used across conditions, with each condition (FRQ and INT) delivered twice, using reverse sessions where the role of the two sounds (standard and deviant) were exchanged. Further details about stimuli and sequences can be found in Lecaignard et al. (2015). Participants were instructed to ignore the sounds and watch a silent movie of their choice with subtitles.

#### Data acquisition

Simultaneous MEG and EEG recordings were carried out in a magnetically shielded room with a whole-head 275-channel gradiometer (CTF-275 by VSM Medtech Inc.) and the CTF-supplied EEG recording system (63 electrodes), respectively. We provide here the aspects of particular relevance for the coregistration of multimodal data. Details regarding the simultaneous MEG and EEG recordings and the experimental setup can be found in (Lecaignard et al., 2015). EEG electrode positions relative to the fiducials were localized using a digitization stylus (Fastrak, Polhemus, Colchester, VT, USA). Special care was taken to minimize head position drifts inside the MEG helmet between sessions. T1-weighted magnetic resonance imaging images (MRIs) of the head were obtained for each subject (Magnetom Sonata 1.5 T, Siemens, Erlangen, Germany). High MRI contrast markers were placed at fiducial locations to facilitate their pointing on MRIs and thereby minimize coregistration errors. Head position relative to the MEG sensors was acquired continuously (continuous sampling at a rate of 150 Hz) using head localization coils placed at fiducial points.

#### Auditory event-related field/potential (ERF/ERP)

MEG evoked responses (2-45 Hz) were computed in exactly the same way as EEG ERPs (Lecaignard et al., 2015), with MEG-specific preprocessings, namely the rejection of data segments corresponding to head movements larger than 15 mm relative to the average position (over the 4 sessions) and to SQUID jumps. Importantly, we only used time epochs that survived the procedures applied for artifact rejection for both modalities. EEG evoked responses were re-referenced to the average of the signal at mastoid electrodes in the current study for compatibility with the forward model. Grand-average responses at gradiometer MLP56 and electrode FCz in condition FRQ and INT are shown in *Figure **3***. Permutation tests (Lecaignard et al., 2015) revealed an early deviance and an MMN in both modalities (EEG, MEG) and both conditions.

**Figure 3.**
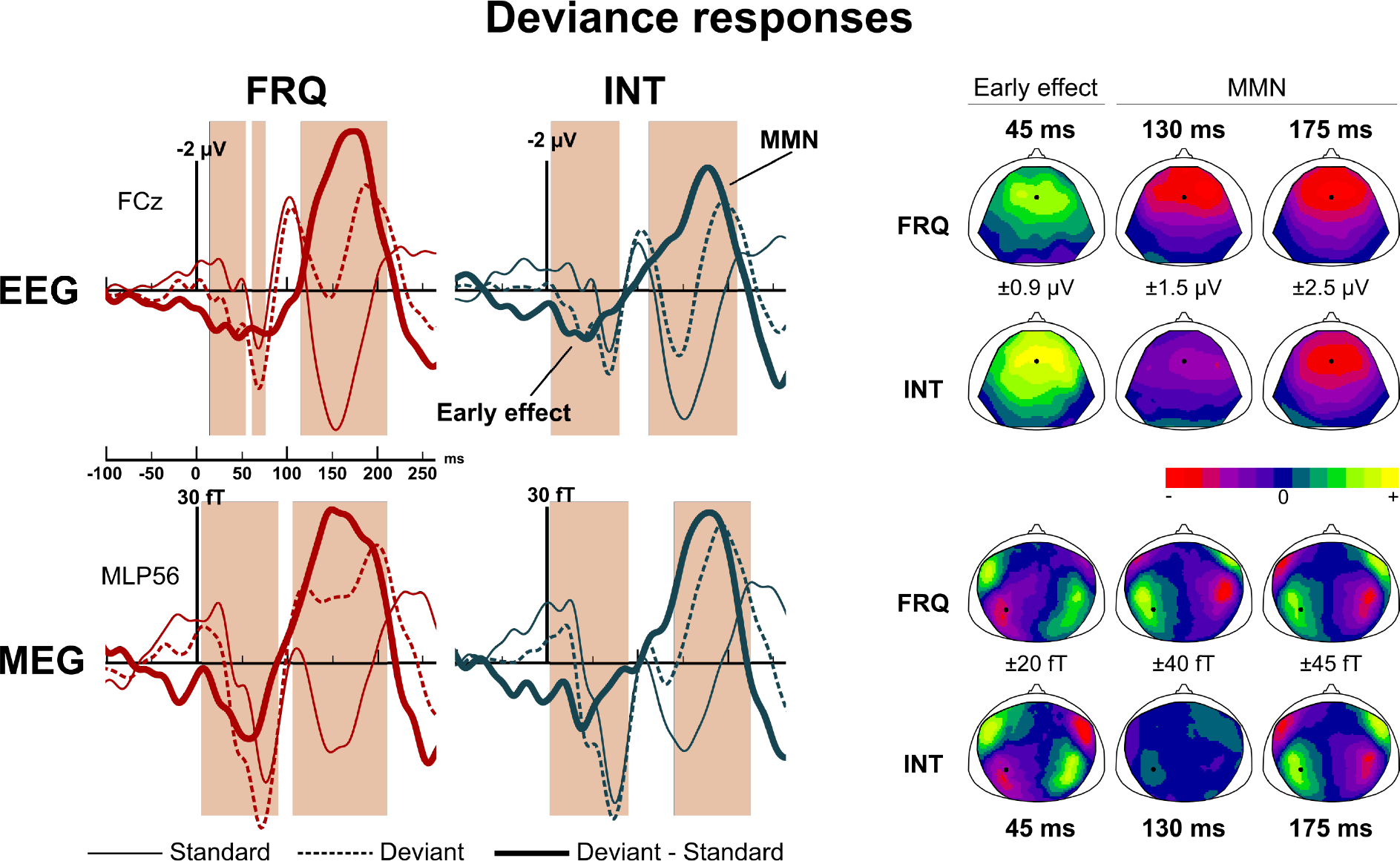
Mismatch ERPs/ERFs. Left panel: auditory evoked responses at electrode FCz (upper row) and gradiometer MLP56 (lower row) for the frequency (left) and intensity (right) conditions. Shaded areas correspond to the time intervals of significant mismatch emergence over all sensors (modality-condition): (EEG-FRQ): [15 55] ms, [65 80] ms, [115 210] ms; (EEG-INT): [5 80] ms, [113 210] ms; (MEG-FRQ): [5 90] ms, [105 210] ms; (MEG-INT): [3 90] ms, [140 225] ms. Right panel: scalp topographies at relevant latencies for the early deviance, the rising edge and the peak of the MMN. Color-scale range is indicated for each map.

#### Data for source reconstruction

We used SPM8 software (Wellcome Department of Imaging Neuroscience, http://www.fil.ion.ucl.ac.uk/spm). Standard and deviant ERFs and ERPs (with averaged mastoid reference) were down-sampled (200Hz) for data reduction. Source reconstructions were estimated for difference responses (deviant-standard) in each condition separately (FRQ, INT) and for each modality (EEG, MEG, Fusion). As sensor-level traces showed a tendency for the intensity MMN to start later than the frequency one, we distinguished the rising edge from the peak of this component to increase the spatial sensitivity of reconstructions. Three time windows were thus considered: from 15 to 75 ms (early deviance effect), from 110 to 150 ms (MMN rising edge), and from 150 to 200 ms (MMN peak). Overall, a total of 18 separate inversions were computed for each of the 20 participants (3 time-windows × 2 conditions × 3 modalities). In addition, our comparative evaluation of separate (EEG, MEG) and fused (MEG-EEG) inversions was applied to the time interval [150,200] ms in both conditions (FRQ, INT). Regarding data normalization, 7 and 21 spatial modes (explaining 99.0% and 99.9% of the group-informed gain matrix variance) were retained for EEG and MEG, respectively. Data reduction using temporal modes was achieved for all inversions. The number of temporal modes allowing for 100.0% of the variance of the spatially projected data to be explained was equal to 6, 4 and 5 for [15, 75] ms, [110, 150] ms and [150,200] ms time intervals, respectively.

#### Statistical analysis on source distributions

We conducted our statistical analyses at the group-level using the recent surface-based approach proposed in SPM12. Posterior estimates of source activity and associated variance at each node of the cortical mesh (the source domain) resulted from posteriors of *J* and *C*_*s*_. The energy of posterior mean was considered for statistical analysis. One-sample t-tests were performed at each node, thresholded at *p* < 0.05 with Family Wise Error (FWE) whole-brain correction. In addition, we imposed the size of subsequent significant clusters to be greater than 20 nodes. Distance between two local maxima within a cluster was constrained to be larger than 5 nodes.

## 3. Results

We first present the comparative evaluation for EEG, MEG and fused inversions that we conducted with FRQ and INT difference responses, at the MMN peak ([150, 200] ms). Second, as multimodal comparison was in favor of fused MEG-EEG inversion, we report the corresponding sources obtained for the time intervals [15, 75] ms, [110, 150] ms and [150,200] ms in the difference responses, in both conditions FRQ and INT, thus applying the current multimodal framework for source reconstruction to the localization of the sources of auditory mismatch responses.

### 3.1. Multimodal evaluation

For each modality, source reconstructions were computed for each subject using difference responses in the time interval [150, 200] ms. Resulting *R* (the percentage of explained variance) in condition FRQ was equal on average to 95.1% (± 2.1), 94.2% (± 2.3) and 93.6% (± 2.6) for EEG, MEG and fused inversions respectively. In condition INT, it was equal on average to 94.7% (± 2.5), 93.8% (± 2.3) and 93.1% (± 2.7) for EEG, MEG and fused inversions respectively. Regarding the contribution of each modality (EEG, MEG) in the case of fused inversion, paired Student’s t-tests were used to compare the estimated values of hyperparameters λ_*e*_ and λ_*m*_. In both condition FRQ and INT, inversions across subjects led to no significant difference between modalities (t(19)=1.30, p=0.21 for FRQ; t(19)=1.98, p=0.06 for INT).

#### Separate and fused MMN source distributions (qualitative comparison)

***Figure 4*** shows the results of the statistical analysis projected on the inflated cortical surface in each modality (EEG, MEG and MEG-EEG) and each condition (FRQ, INT). In both conditions, EEG and MEG inversions led to different (but not inconsistent) reconstructed activity, and more focal clusters were found with fused inversion. Precisely,

- In condition FRQ, EEG inversion revealed bilateral activity in the anterior part of the supratemporal plane and in the lower bank of the posterior STG. No frontal area was found significant. MEG inversion indicated a large cluster in the supratemporal plane (number of nodes k > 120) expanding from the lateral part of HG through the Planum Polare (PP) in both hemispheres. A bilateral frontal area was located in the posterior IFG. The fused distribution comprised smaller supratemporal clusters (right: a single cluster (k=92) including the lateral part of HG and PP; left: separate clusters for HG (k=55) and PP (k=25)), and bilateral clusters similar to MEG ones in the frontal lobe.
- In condition INT, the EEG solution indicated bilateral activity in the posterior STG and the intraparietal sulcus (IPS). There was a similar distribution to condition FRQ with MEG. Fused inversion gave largest contributions in the lateral part of HG in both hemispheres, but also right clusters located in posterior IFG, posterior STG and in the inferior temporal gyrus (ITG).

**Figure 4.**
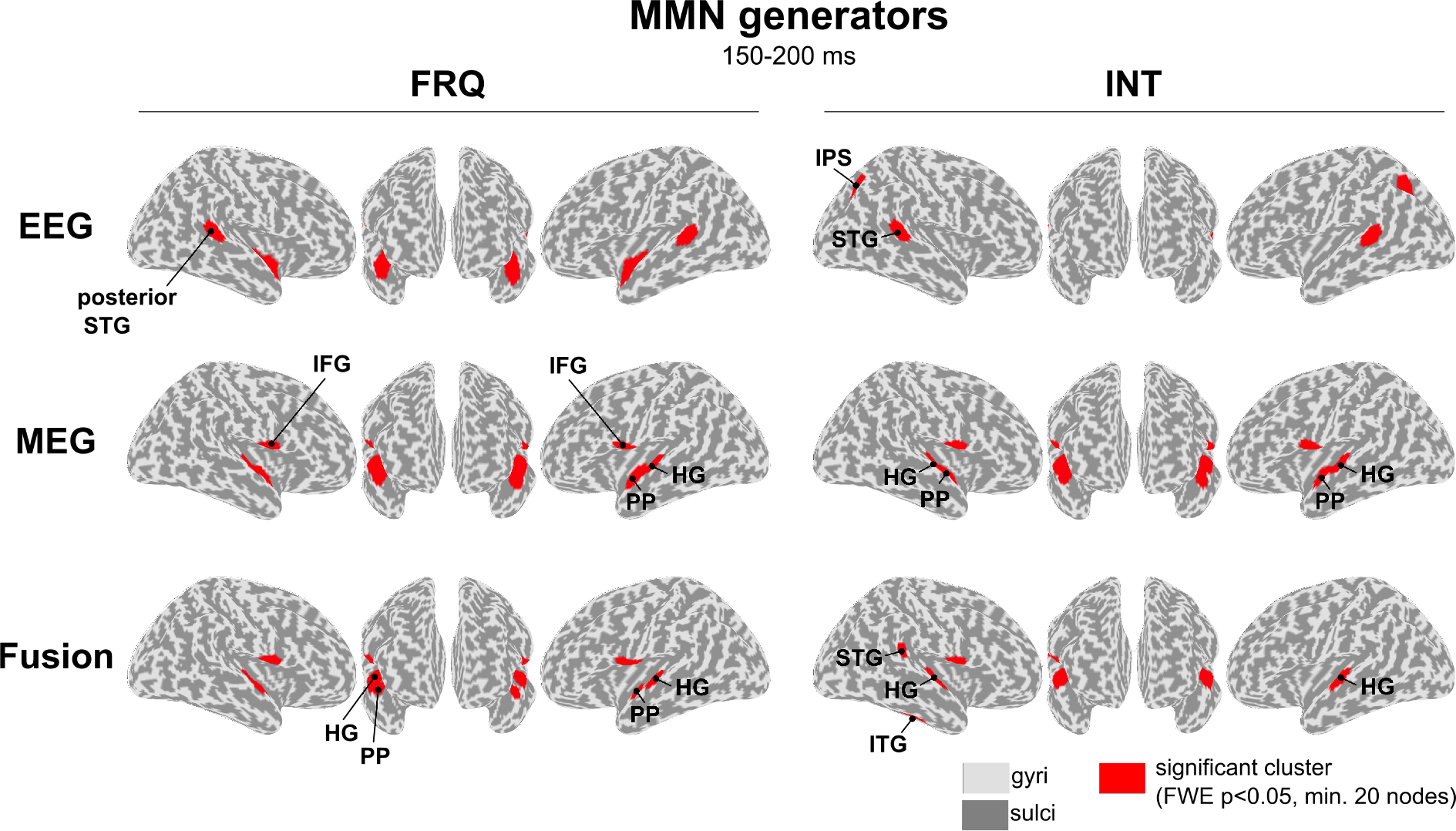
Comparison of the EEG, MEG and fused MSP inversions corresponding to the MMN peak for frequency (left panel) and intensity (right panel) deviance. Red clusters indicate the significant source activity over the group (N=20) projected on the inflated cortical surface (HG=Heschl’s gyrus; STG= superior temporal gyrus; P=planum polare; IFG=inferior frontal gyrus; IPS= inferior parietal sulcus; ITG=inferior temporal gyrus).

#### Group prior models (***Figure 5***)

Group priors obtained in condition FRQ vary across modalities, with EEG priors strongly diverging from MEG and Fusion ones. Precisely, EEG upweights bilateral priors in posterior STG and the anterior temporal lobe. In contrast, MEG and Fusion group priors appear similar, upweighting bilateral posterior IFG and the supratemporal plane (including HG and PP). In the following, we assume that MEG and Fusion group prior models consist in *close* models, whereas EEG and MEG ones, and EEG and Fusion ones are *distant* over model space. Contrary to condition FRQ, group priors in condition INT are different across all the three modalities. EEG priors were located bilaterally in posterior STG, ITG and IPS. Remarkably, MEG and Fusion both provided bilateral priors in posterior IFG priors, but strongly differed in the supratemporal plane, with MEG involving lateral HG and PP while Fusion focused on HG only. Like in condition FRQ, models from EEG and MEG, and from EEG and Fusion were assumed to be *distant* over model space. Regarding MEG and Fusion models, difference related to PP led us to assume that they were more distant in condition INT than in condition FRQ. In both conditions and for all modalities, less restrictive priors (smaller cluster size and/or larger variance) were also found that we do not report here for they did not survive any individual inversion.

**Figure 5.**
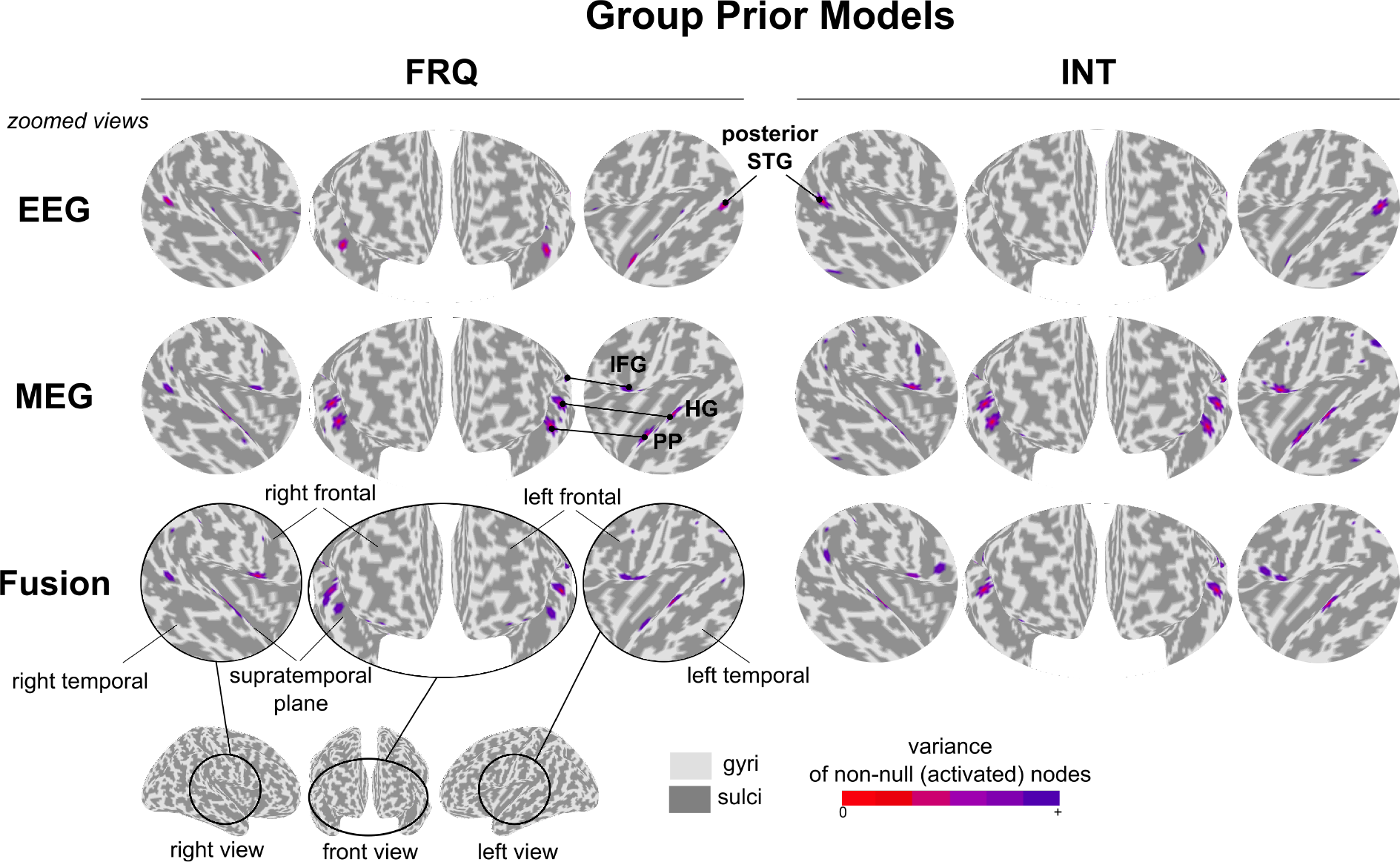
Comparison of group prior models obtained for EEG, MEG and fusion (in separate rows), with the frequency (left panel) and intensity (right panel) deviances. For each modality and each condition, three zoomed views (with relation to the global view indicated at the bottom left) indicate the result of MSP inversion performed at the group level (first step of group-level inference), hence reflecting the source distribution common to the group.

#### Bayesian Model Comparison (***Figure 6***)

Three BMC per condition were conducted based on the approximated model evidences inferred from the nine cross-modal inversions. In condition FRQ, no difference between modalities was measured: model 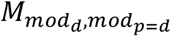 was each time selected as the winning model. Precisely, BMC gave the following model exceedance probabilities: *p*(*M*_*e,e*_|*Y*_*e*_) = 1.00, *p*(*M*_*m,m*_|*Y*_*m*_)=0.97 and *p*(*M*_*f,f*_|*Y*_*f*_)=0.88. In condition INT, model 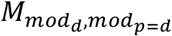 won in the case of EEG and MEG-EEG inversions, with *p*(*M*_*e,e*_|*Y*_*e*_) =1.00 and *p*(*M*_*f,f*_|*Y*_*f*_) =0.97, but BMC was inconclusive for MEG inversion, (*p*(*M*_*m,m*_|*Y*_*m*_) = 0.52 and *p*(*M*_*m,f*_|*Y*_*m*_) =0.48, respectively). In conclusion, in all cases but MEG inversion in condition INT, model separability was always evidenced with each time the preferred model corresponding to the group-level solution obtained with the modality used for inversion 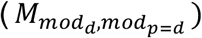. In the case of MEG in condition INT, fused priors were found performing as well as MEG ones, despite their difference related to the implication of PP. The opposite was not observed: in fused inversion, BMC clearly decided in favor of Fusion priors. This asymmetrical result demonstrates larger model separability in Fusion than in MEG.

**Figure 6.**
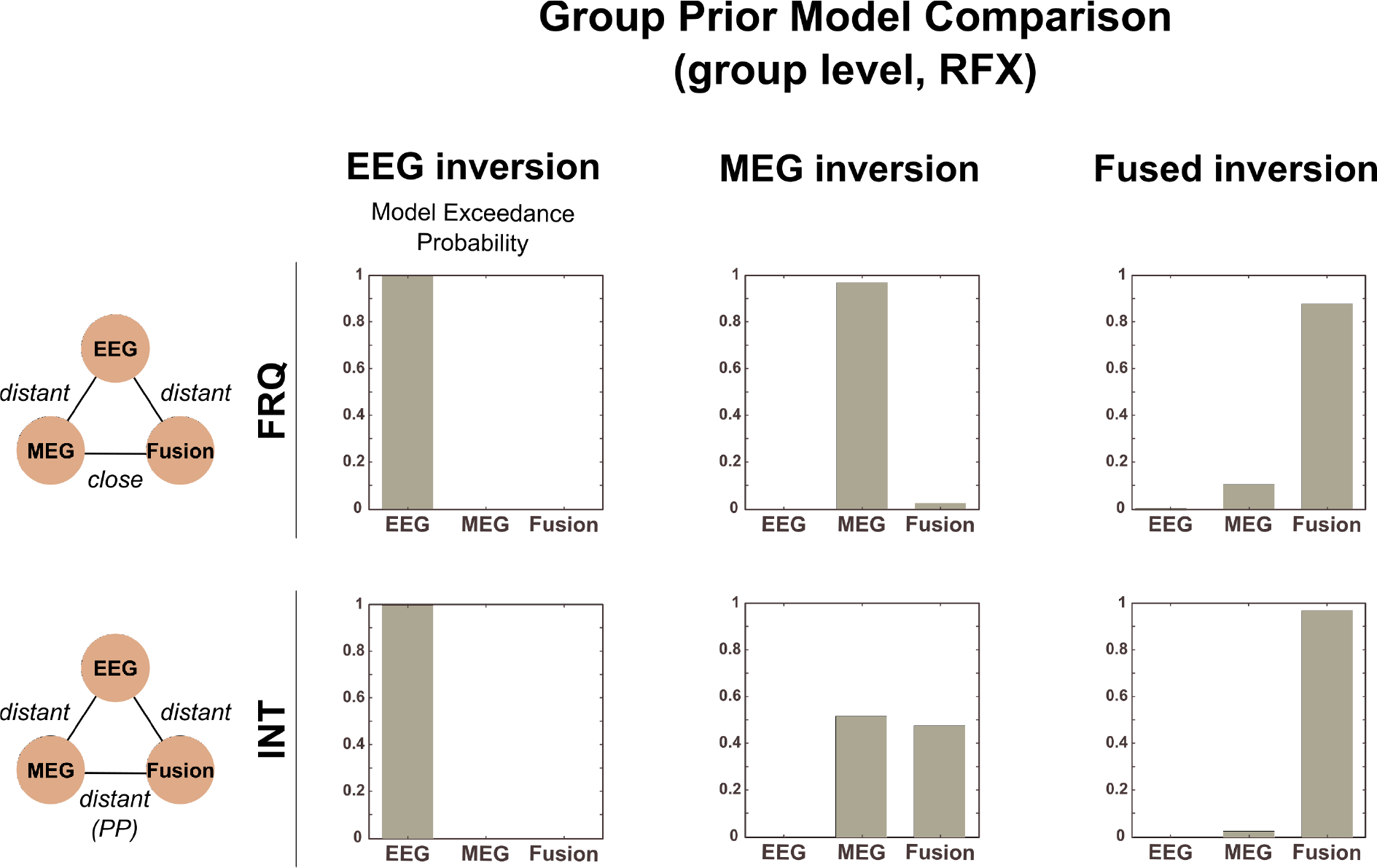
Bayesian model comparison of group prior models within each modality (in separate columns) for the frequency (upper row) and intensity (lower row) deviances. Each graph displays model exceedance probabilities for each prior model. Diagrams on the left summarize the distance between models that we classified into close or distant models. This highlights the fact that the lower model separability of MEG than fusion could be revealed in the case of distant models, as was the case for INT.

#### Individual free energy difference (***Figure 7***)

This second analysis accounting for within-subject variability enabled us to identify even more subtle patterns across modalities. The key points to take from ***Figure 7*** are:

- EEG exhibited a poor ability to separate models: EEG inversion was found to perform as well with EEG group priors than with MEG and fused ones. In addition, EEG inversion at the group level provided solutions (the group priors) that were rejected by other modalities, suggesting that priors derived from EEG inversions were too poorly informed to be compatible with MEG data.
- Regarding MEG and Fusion performances: in the (easy) case of clearly distinct models (EEG vs. MEG priors, EEG vs. Fusion priors, in all conditions), individual results indicate that both MEG and Fusion could resolve models. In the intermediate case (MEG vs. Fusion priors, condition INT) and in the case of close priors (MEG vs. Fusion priors, condition FRQ), differences between modalities can be reported: MEG could not conclude in favor of a model in most subjects (14/20 inconclusive in both conditions), whereas Fusion could (12/20 and 14/20 conclusive in FRQ and INT, respectively) and selected fused priors (9/12 and 12/14, respectively).

**Figure 7.**
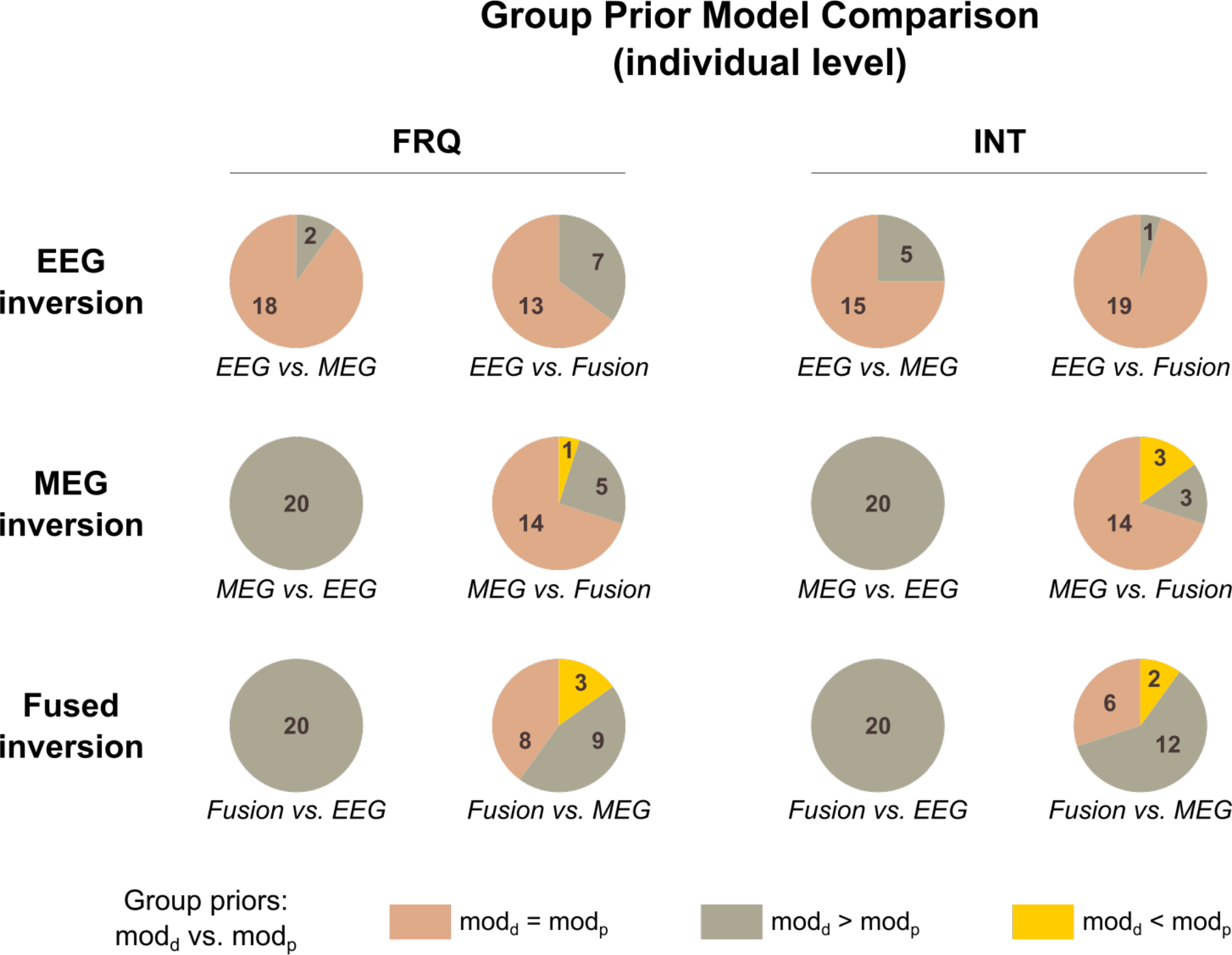
Model comparison based on individual free energy differences computed for each modality (in separate rows) and for each condition (in separate columns). The two diagrams provided for each modality and each condition summarize the free energy differences (over the group) between this modality and the two others, separately, as indicated below each graph. Pink areas indicate the proportion of subjects (with exact number indicated) for whom model comparison concludes in favor of equal plausibility (absolute difference ≤ 3). Grey and yellow areas indicate the proportion of subjects in favor of group priors from the concerned modality (diff. > 3) and from the other one (diff. < −3), respectively. In short, the smaller the pink area, the larger model separability. *mod*_*d*_ and *mod*_*p*_ refer to the modality used for data inversion and for group priors, respectively.

#### Summary

Reconstructions of the sources of the frequency and intensity MMN were performed using EEG, MEG and MEG-EEG inversions. Source distributions all provided a very good fit of data and in fused inversion EEG and MEG contributed equally to the inversion process. Our evaluation approach relying on group prior model comparison succeeded at quantifying the performance of each modality for the reconstruction of empirical data. Precisely, we found larger performances for fused inversion to resolve group-level informed distributions. In the following section, we therefore present the deviance-related source reconstructions obtained only with this modality.

### 3.2. Fused MEG-EEG sources for auditory mismatch responses

*Figure **8*** shows the results obtained for each deviance type and each time interval with fused inversion. Cluster sizes and peak location in MNI space for each local maxima for significant activated areas are summarized in ***Table 1*** for condition FRQ, and ***Table 2*** for condition INT.

**Figure 8.**
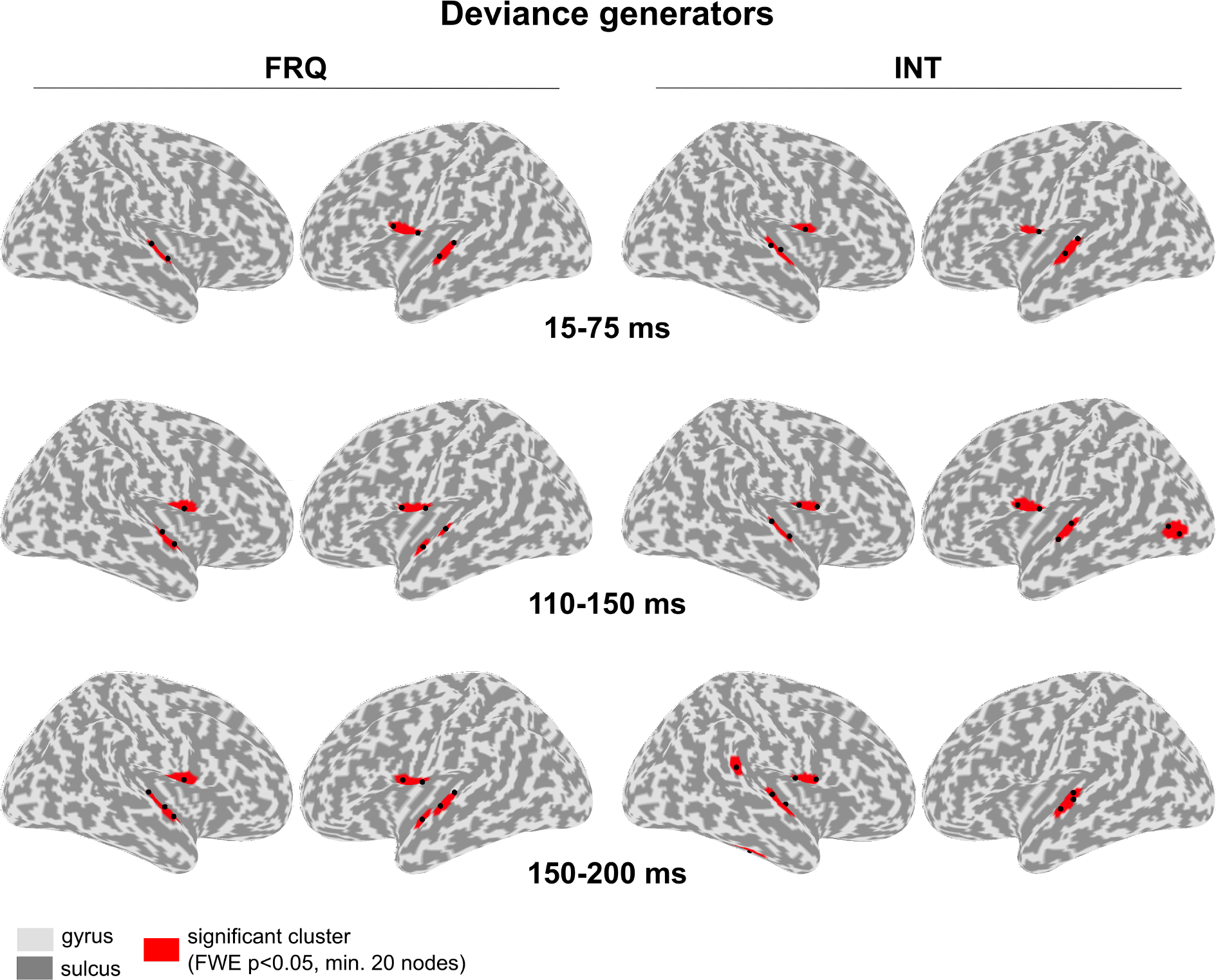
Deviance generators (fused MSP reconstruction). Significant clusters (red) are displayed on the inflated cortical surface (right and left views) for each time interval (rows) and each condition (frequency =left panel, intensity=right panel). Black dots indicate the local maxima within each cluster (with a minimum distance of 5 adjacent nodes). MNI coordinates are provided in Table 1 (frequency) and Table 2 (intensity).

**Table 1.**
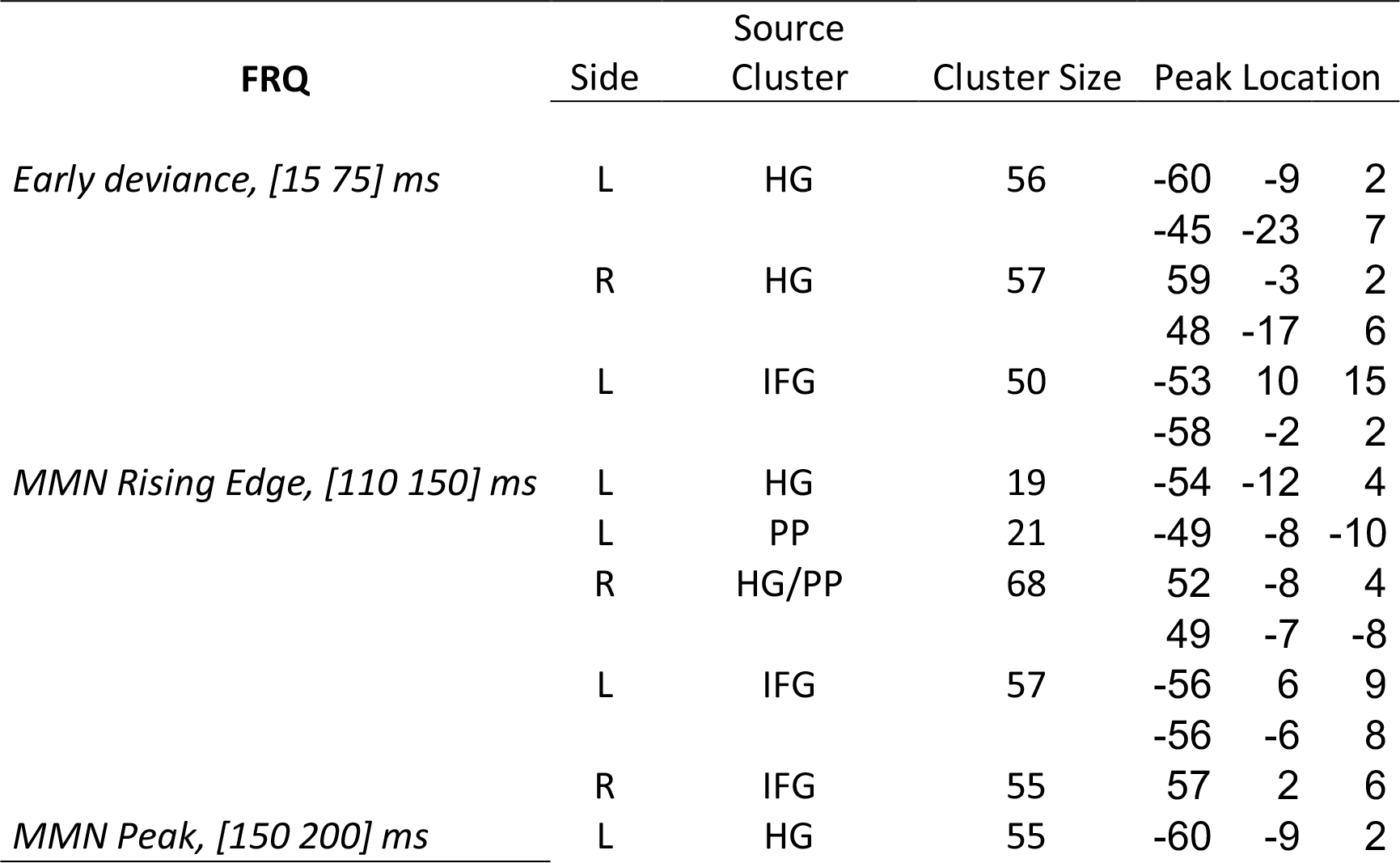
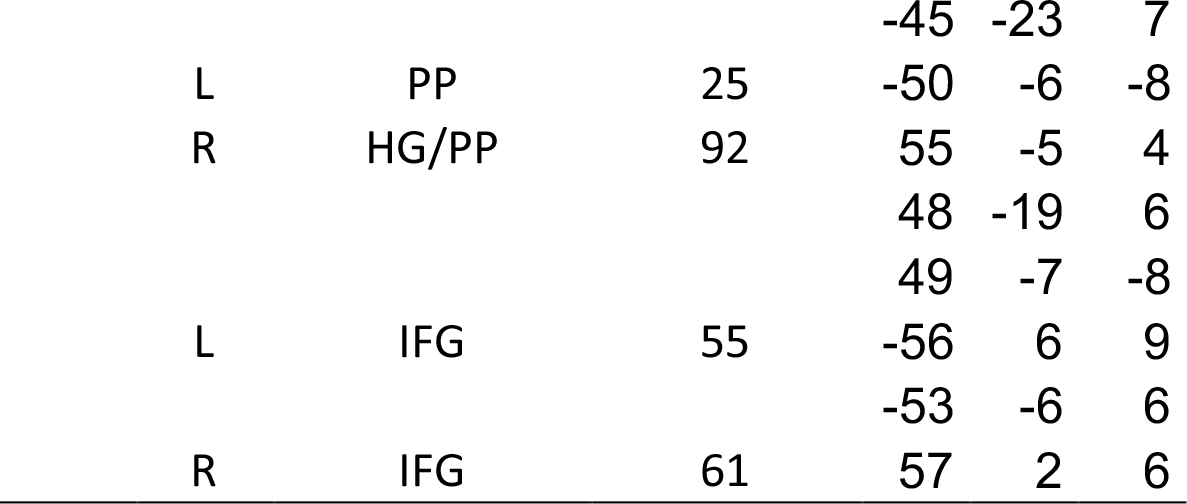
Results of MSP inversion for frequency deviance with fused inversion. Condition INT.

**Table 2.**
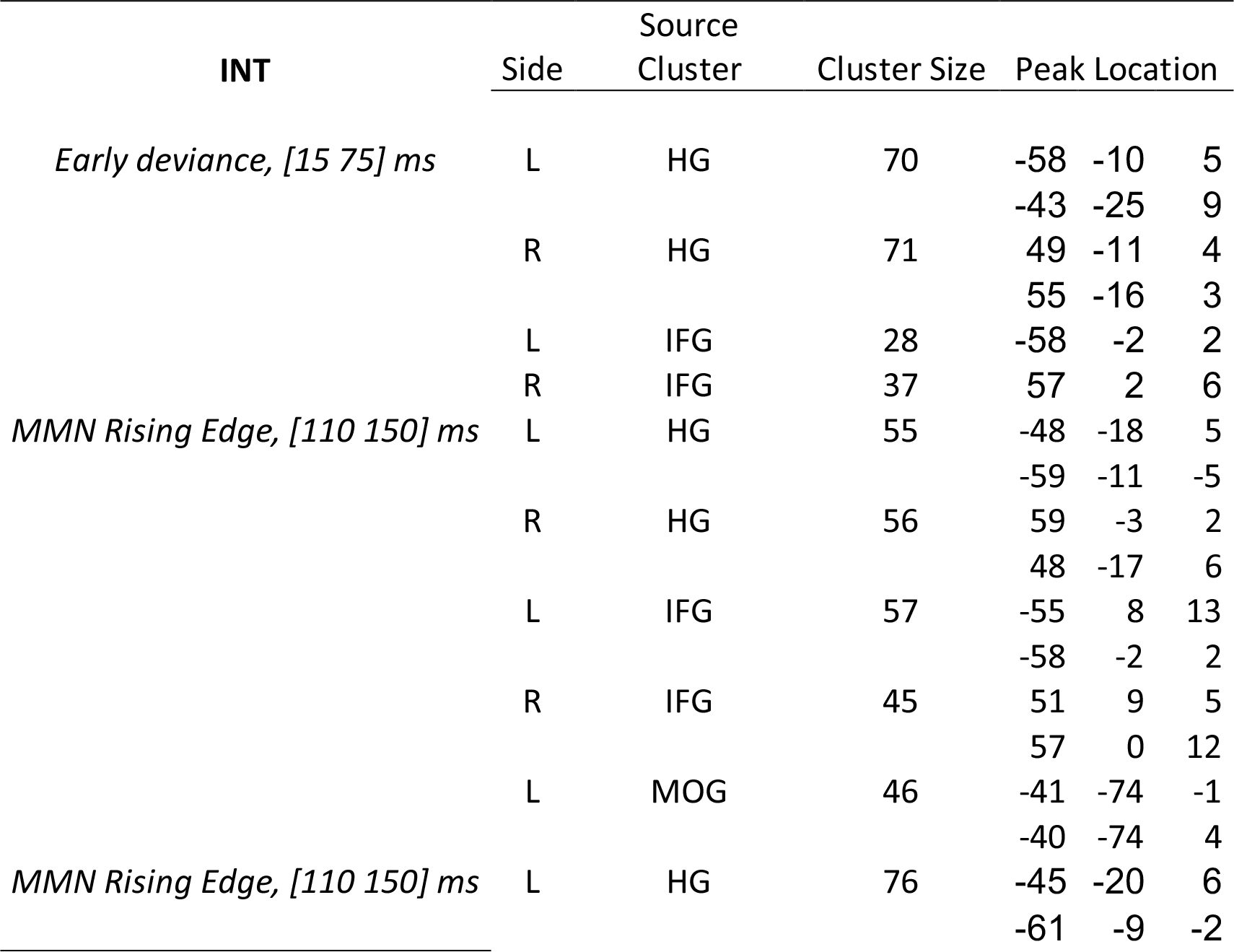
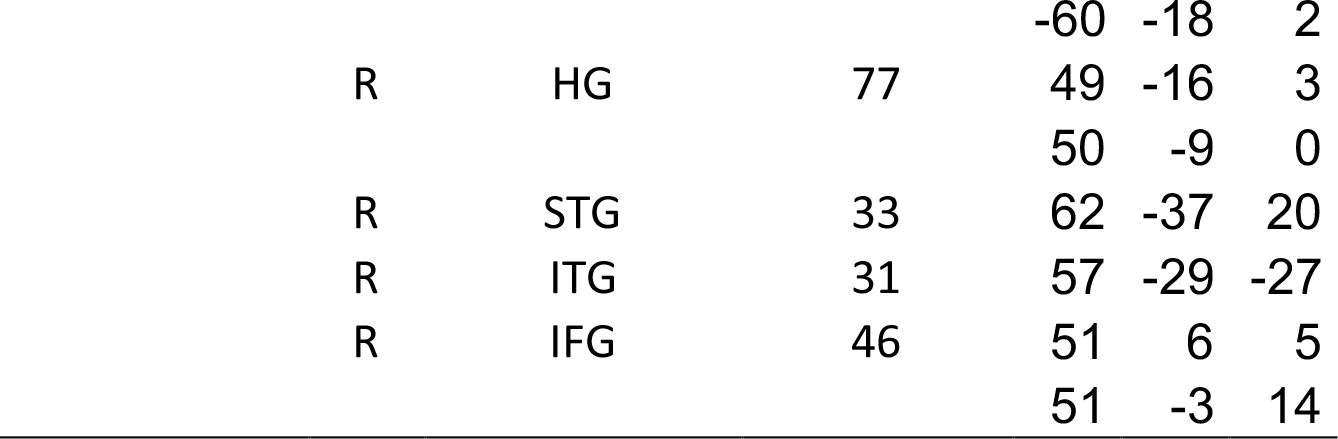
Results of MSP inversion for intensity deviance with fused inversion

#### Condition FRQ

Reconstructions of deviance generators within time windows [15, 75] ms, [110, 150] ms and [150,200] ms were performed with resulting *R* equal on average to 90.7% (± 4.8), 92.3% (± 4.4) and 93.6% (± 2.6), respectively. Early-deviance effect ([15, 75] ms) was found to involve HG in both hemispheres and left posterior IFG. Following this, reconstruction of the rising edge of the MMN ([110, 150] ms) indicated supratemporal activity in HG and PP, within a large cluster in the right hemisphere (comprising two local maxima), and separated in two distinct clusters in the left hemisphere (with HG cluster being smaller). Significant activity was also found in bilateral posterior IFG. Finally, as described in previous section, the peak of the MMN ([150,200] ms) was associated with activity in both hemispheres peaking in HG, PP and posterior frontal IFG. The total number of significant nodes within bilateral supratemporal planes was larger for the peak than for the rising edge of the MMN (178 and 108 respectively), while it remained constant within IFG (116 and 112 respectively).

#### Condition INT

*R* was equal on average to 91.4% (± 5.2), 90.5% (±5.4) and 93.1% (±2.7) for the reconstructions within time windows [15, 75] ms, [110, 150] ms and [150,200] ms, respectively. Within the early-deviance window ([15, 75] ms), activity was mostly found in bilateral HG but was also located in posterior IFG. Reconstructions within [110, 150] ms produced significant clusters in bilateral HG and posterior IFG. In addition, there was a contribution from left middle occipital gyrus (MOG). Finally, sources in HG and posterior IFG were observed in both hemispheres for the MMN peak reconstruction ([150,200] ms). Smaller clusters were found in ITG and posterior STG in the right hemisphere. With the thresholds chosen in the current study, no contribution of PP could be reported at any latency.

#### Summary

The fused reconstructions of deviance responses observed in ERP/ERF revealed a bilateral fronto-temporal network in both conditions (FRQ, INT). Temporal activity was clustered in the supratemporal plane, where fused inversion improved the spatio-temporal description of deviance-related activity. In particular, fused inversion could separate HG and PP clusters spatially, but also temporally as PP contribution varies over time and across conditions. Frontal contributions could be recovered in both conditions as soon as the early deviance window.

## 4. Discussion

In the present study, we propose a general approach to tackle the long-standing issue of quantifying the gain of EEG-MEG fusion in empirical source reconstruction. This was achieved here by no longer addressing the intractable problem of localization performances but by considering the spatial separability of modalities that can be formally assessed in a Bayesian framework. State-of-the-art methodologies employed throughout the study encompass an advanced realistic forward model, Bayesian inverse methods with group-level inversion and fused EEG-MEG inversion, coupled with surface-based statistical tools. Evaluation applied to the reconstruction of MMN sources revealed a larger resolution with fused inversion, as predicted by the existing simulation-based literature. Fused inversion applied to early and late deviance responses resulted in a fronto-temporal network consistent with EEG and MEG alone existing findings, but described here to our knowledge with a spatio-temporal precision not hitherto attained.

### A general method to compare modalities for distributed source reconstruction

The originality of the proposed procedure pertains to its suitability for empirical data, without the need for establishing *a priori* a true distribution to refer to. It fully exploits advantages of the Bayesian inversion framework, namely the acknowledged Bayesian model selection and recent group-level inference. In this way, we derived an easy-to-achieve comparison tool, estimating the capacity of each modality to resolve source distributions. Model space for such priors is infinite and was restricted here to three distributions in particular: the group priors (or equivalently, the source covariance common to all subjects) inferred for each modality. Their great relevance for the comparison scheme comes from the fact that they reflect the information gathered by each modality over the group of subjects. Application to auditory responses indicates that each modality selected its own priors (but MEG in condition INT). Following this, individual inspection revealed however the lack of information in EEG data that prevented to disentangle source distributions, whereas MEG and most clearly fusion were sufficiently informed to do so. Looking closely at the differences across the two conditions (FRQ, INT) underlines the importance of the distance between models to refine the evaluation: the large (PP-related) difference between MEG and fusion priors in condition INT revealed the larger separability of fused inversion that was not observable with condition FRQ. Going further, parameterizing distance over model space using synthetic priors could provide a quantitative tool to estimate the spatial resolution of each modality. Importantly, our approach illustrates the power and the flexibility of a statistical framework to test precise hypotheses that, in the present case, allowed us to provide practical guidelines to improve model inversion (namely, we recommend fused inversion to model auditory cortical activations). Bayesian model comparison has already been employed to improve forward modeling (Henson et al., 2009a; Strobbe et al., 2014). In the same vein, one could extend our approach to virtual modalities composed of subsets of sensors to identify the most informed ones (according to our criterion of model separability) to enter subsequent analysis. Sensor selection (in EEG in particular) is an important practical aspect to consider when designing a new experiment.

### Fused inversion has a larger spatial model resolution

Application of our evaluation procedure to auditory mismatch responses (in two separate conditions) indicated a larger ability of fused inversion to separate spatial distributions and to select the fused one. Fused priors (reflecting the source distribution common to the group) appear sufficiently informed to improve EEG, MEG and fusion individual inversion. Of course, generalizability of the present results to other brain activations should be evaluated, that could further help to characterize the spatial complementarity of EEG and MEG recordings. Still, our findings are definitely in accordance with expectations from the simulation-based literature and constitute robust empirical evidence resting on a sizeable group of subjects (N=20) and a quantitative procedure. Importantly, no significant difference between the measurement noise estimate (hyperparameter λ_*n*_) obtained for the two modalities in the case of fused inversion led us to assume that both modalities equally contributed to the inversion. This is an important control that Bayesian framework for inversion provides (Henson et al., 2009b). It strongly supports the great performances of fused inversion being the result of the complementary information gathered by each modalities rather than one modality prevailing the other. We know that EEG and MEG data comprise a mixture of neuronal contributions from various origins, and that they capture different aspects of the same underlying biophysical activity (Lopes da Silva, 2013). An illustration was given here where at the scalp level, there were significant differences between conditions at the latency of the MMN in MEG that were not visible with EEG (statistical analysis not shown in the present report). At the source level, it is well known that EEG and MEG do not have the same sensibility to various assumptions embedded in forward modeling (Lecaignard and Mattout, 2015). They possibly explain the observed differences across unimodal inversions within the IFG (further discussed below) and temporal regions. Regarding more precisely the latter, current MEG sources in the primary auditory cortex corroborate the acknowledged potential of MEG to resolve temporal lobe activity (see for instance early ECD findings in Alho, 1995). Yet, only fused inversion succeeded in revealing subtle patterns within the supratemporal plane with modulations over the temporal dynamics and over deviance features. This highlights the importance to include EEG information to improve MEG spatial resolution in the particular case of supra-temporal activations.

### Mismatch sources with fused inversion

The comparative analysis performed at the peak of the MMN strongly encouraged us to merge EEG and MEG data to finely characterize other mismatch sources, as never done before. The main finding of this subsequent analysis is the identification of a bilateral fronto-temporal network at play during early and late deviance responses and for both conditions (FRQ, INT). The MMN result (including the rising edge and the peak of the MMN) is totally consistent with the existing literature. Perhaps the most striking point is that expected contributions (frontal and temporal sources, bilaterally) could be all identified at once, which is far not so common. The spatial specificity of the network (and in the supra-temporal plane in particular) is also noticeable in comparison to the large cluster sizes often reported. In fact, the present findings are comparable to fMRI results, likewise those reported in the study of Schönwiesner and collaborators (2007). From a qualitative point of view, fused inversion could thus reach the spatial resolution of fMRI (at least in the temporal lobe), which made possible to reveal distinct spatial patterns across specific time intervals in the first 200 ms of auditory processing (which is obviously not feasible with slow metabolic imaging). In particular, we could observe in the supratemporal plane a posterior to anterior progression (from HG to PP) between the rising edge and the MMN peak for the frequency condition, that appears in keeping with several studies that explored the N1 and the MMN generators (Recasens et al., 2014a; Scherg et al., 1989). Comparison of frequency and intensity distributions (although beyond the scope of the study) also shows subtle spatio-temporal patterns: similar activations at early latency are followed by differences within the supratemporal plane and frontal regions during the MMN. This supports different sensory processes at the MMN latency, as proposed by early ECD studies conducted with EEG (Giard et al., 1995) and MEG (Levänen et al., 1996). Regarding early deviance generators, temporal activity was clearly circumscribed within bilateral Heschl’s gyrus for both deviance features. This is totally consistent with MLR findings from intracranial recordings studies (Liégeois-Chauvel et al., 1994; Pantev et al., 1995; Yvert et al., 2002). Recent MEG findings also reported temporal contributions including HG, in the right hemisphere (Recasens et al., 2014a) and bilaterally (Recasens et al., 2014b). Crucially, a major difference with these studies pertains to frontal sources that we were able to recover. Under the assumption of a hierarchical organization for deviance processing that could unfold from subcortical areas to higher cognitive cortical levels (Escera and Malmierca, 2014), such frontal contribution as soon as these early latency has now become highly expected.

### Limitations of mismatch findings

Unexpected findings reported here should however be discussed. First concerns the failure to identify any frontal contribution with EEG at the MMN peak. Deep inspection of the MMN literature reveals that it is not straightforward to locate the IFG with EEG responses, unless considering specific priors with discrete ECD models (Jemel et al., 2002; MacLean et al., 2015; Rissling et al., 2014). With distributed source localization, two recent reports of an IFG contribution can be cited (Fulham et al., 2014; in a language study: Hanna, 2014). In our case, it is likely that inferior frontal activations were less plausible (possibly weaker) than supra-temporal ones, and as such they have been canceled out by MSP, which implements the principle of sparsity to activated sources. It should also be noted that few MEG studies also succeeded in localizing these regions (Lappe et al., 2013b; Recasens et al., 2015). Another unexpected result pertains to the contribution of the left middle occipital gyrus and the right inferior temporal gyrus for intensity deviance with fused inversion. It is worth recalling that the intensity MMN was not significant at the scalp-level over the time interval from 100 to 150 ms. We therefore assume that these sources constitute false positive (deriving from a convergence into local minima). Finally, frontal contributions were located in the very posterior part of the IFG, mirroring supratemporal regions. The fact that we observed activations in IFG but not in PP (rising edge of the MMN, INT) and in HG but not in IFG (peak of the MMN, INT) allows to reject the hypothesis of two mis-localized and correlated clusters of opposite sign. All these considerations constitute however a reminder of the ill-posed nature of the source reconstruction problem, that will always remain whatever the advanced methodologies integrated in the inversion framework.

## 5. Conclusion

This paper develops an evaluation procedure to test quantitatively the gain of fusing EEG and MEG data for distributed source localization. Critically, it consists in a general approach that applies to empirical data. It thus paves the way to go beyond stimulations to test the better performances of fusion predicted by biophysical and information theory principles. From a practical point of view, it also appears convenient for the experimenter to quantify the gain of fusion for particular brain responses of interest. In the present case of auditory responses with recent methods in distributed source modeling, fusion was found to outperform EEG and MEG alone, as expected, and can now be formally highly advised in subsequent auditory studies. We identified a bilateral fronto-temporal network for both frequency and intensity deviance responses that conforms the existing mismatch literature. Promisingly, the spatial resolution reached with fused inversion allowed a detailed spatio-temporal description within the supratemporal plane. These findings should however be balanced against the experimental cost of simultaneous EEG-MEG acquisitions that remain somewhat less straightforward that unimodal ones. Still, they should be considered as an attractive and powerful option that we recommend, particularly in the case of auditory studies. As a result, the refined auditory network achieved here represents a crucial step to further address auditory processing using mismatch responses, be it at the neurophysiological or the cognitive levels.

